# LSD1/KDM1A is essential for neural stem cell differentiation in mice

**DOI:** 10.1101/2023.12.02.569711

**Authors:** E.C. Falkenberry, M. Reeves, A. Scott, D.A Myrick, C. Fallini, G.J. Bassell, D.J. Katz

## Abstract

The proper regulation of neural stem cell differentiation is required for the proper specification of the central nervous system. Here we investigated the function of the H3K4me1/2 demethylase LSD1/KDM1A during neural stem differentiation in mice. Conditional deletion of LSD1 in *nestin -* positive neural stem cells results in 100% perinatal lethality after birth with severe motor coordination deficits, retarded growth and defects in brain morphology. Despite these severe defects, motor neuron progenitors and the initial motor neuron population are specified normally and motor neurons with normal morphology can be cultured from these mice *in vitro*. However, motor neurons cultured from mice lacking LSD1 in neural stem cells continue to inappropriately maintain critical neural stem cell proteins. Taken together these results suggest that, as in other mouse stem cell populations, LSD1 is required to deactivate the stem cell program to enable normal neural stem cell differentiation. However, unlike in other mouse stem cell populations, the inappropriate maintenance of the stem cell program during neural stem cell differentiation may compromise neuronal function rather than neuronal specification.

## Introduction

Stem cell (SC) populations have the unique ability to self-renew as well as differentiate into various cell types (Zakrzewski et al., 2019). Lysine 4 of histone H3 (H3K4) methylation is typically associated with active gene expression and has been shown to function as an epigenetic memory, maintaining transcriptional states over time and through cell divisions (Mito et al., 2005; Muramoto et al.; Ng et al., 2003). This transcriptional memory function is important in the maintenance of cell fates, including stem cell programs (Liang & Zhang, 2013). As a result, transcriptional memory needs to be erased in order for proper differentiation to occur.

LSD1/KDM1A (hereafter referred to as LSD1) is an H3K4 demethylase that uses an amine oxidase reaction to specifically remove mono- and di-methylation (me1/2) (Shi et al., 2004). LSD1 plays an essential role in mouse embryonic stem cells (mESCs), decommissioning the active enhancer modification H3K4me1 (Whyte et al., 2012). Without LSD1, H3K4 methylation is not removed at critical SC gene promoters and enhancers, resulting in the inappropriate expression of SC genes after mESC differentiation. Thus, the removal of H3K4 methylation by LSD1 is required for changes in cell fate.

The function of LSD1 in decommissioning enhancers is also required *in vivo* in hematopoietic SCs (Kerenyi et al., 2013; Saleque et al., 2007), trophoblast SCs (Zhu et al., 2014), testis SCs (Lambrot et al., 2015; Myrick et al., 2017), naïve B cells (Haines et al., 2018), satellite SCs (Tosic et al., 2018) and human embryonic SCs (Adamo et al., 2011). The knockout of LSD1 in all of these cell types has been shown to significantly reduce the ability of those SCs to properly undergo differentiation. LSD1 may also function mouse neural stem cell (NSC) differentiation. NSCs give rise to the central nervous system during embryonic development.

LSD1 is expressed in NSCs, and that expression decreases with differentiation, which coincides with elevated H3K4me2 (Sun et al., 2010). Sun et al. also showed that chemical inhibition and siRNA knockdown in NSCs led to inhibition of NSC proliferation in cell culture and in the hippocampus of adult mice (Sun et al., 2010). Furthermore, knockdown of LSD1 inhibits human NSC differentiation (Hirano & Namihira, 2016). This role of LSD1 in human NSC differentiation may be mediated by the microRNA miR-137 (Sun et al., 2011). Taken together, these data indicate a requirement for LSD1 during NSC differentiation. However, the function of *Lsd1* in NSCs differentiation during mouse embryonic development has not yet been examined.

To address this gap, our lab utilized *Nestin-Cre* to delete *Lsd1* in NSCs during embryonic development. The *Nestin-Cre* transgene is strongly expressed in neuronal tissues from embryonic day 10.5 (e10.5) through adulthood, during the time period that NSCs begin to proliferate into various progenitor cell populations (Tronche et al., 1999). The mice in which *Lsd1* is deleted in *Nestin* positive NSCs will hereafter be referred to as *Lsd1*^*NSC*^. While *Lsd1*^*NSC*^ animals make it out to birth, 100% of them die before weaning. Prior to death, *Lsd1*^*NSC*^ mice show a dramatic reduction in size compared to controls and exhibit motor defects. These defects are associated with abnormal brain morphology. Additionally, when motor neurons are cultured from *Lsd1*^*NSC*^ mice, we find that they continue to inappropriately maintain critical SC proteins, including NESTIN and SOX2. These data indicate that LSD1 functions in the proper differentiation of embryonic NSCs, analogous to LSD1’s function in other stem cell populations.

## Materials and Methods

### Mouse husbandry and genotyping

The following mouse strains were used: *Lsd1* floxed mice MGI: 3711205 (Wang et al., 2007) and *Nestin-Cre* MGI: 4412413 (Tronche et al., 1999). Primers for Cre F: GAACCTGATGGACATGTTCAGG, Cre R: AGTGCGTTCGAACGCTAGAGCCTGT, Cre ctrl F: TTACGTCCATCGTGG ACAGC Cre ctrl R: TGGGCTGGGTGTTAGCCTTA. If Cre+, this results in a 302 bp product, and Cre ctrl F/R primers are an internal control that yields a 250bp product. Primers for Lsd1 forward (F): GCACCAACACTAAAGAGTATCC, Lsd1 reverse (R): CCACAGAACTTCAAATTACTAAT. A wild type allele of *Lsd1* results in a 720 base pair (bp) product, the floxed allele is 480 bp, and the deleted allele is 280 bp. All mouse work was performed under protocols approved by the Emory University Institutional Animal Care and Use Committee.

### Histological methods

For immunofluorescence on embryos, we set up timed matings. Pregnant females were sacrificed using cervical dislocation and embryos were fixed for 1-2 hours at 4°C in 4% paraformaldehyde, followed by a 2 hour PBS wash and then transferred to 30% sucrose overnight at 4°C. The tissue was then embedded in O.C.T. compound (Tissue-Tek) and stored at -80°C. 7-10μm sections were incubated with primary antibody in wash solution (1% heat-inactivated goat serum, 0.5% Triton X-100 in PBS) overnight at 4°C in a humidified chamber. Slides were incubated in secondary antibody (1:500 in wash solution) at room temperature for 2 hours in humidified chamber. Antibodies used include: KDM1A (1:250, Abcam ab17721), Cre (1:200 Sigma-Aldrich C7988), HB9 (1:10 DSHB81.5C10), and Olig2 (1:250 Abcam ab9610).

For immunofluorescence on cell, cells were fixed at 3 DIV with 4% paraformaldehyde in PBS and stained with anti-tau (1:5000, Aves), anti-Nestin (1:200, Abcam), and anti-SOX2 (1:200, Epitomics) antibodies overnight at 4°C. Cy3-, Cy2- or Cy5-conjugated secondary antibodies (Jackson Immunoresearch) were incubated for 1 hour at room temperature.

Hematoxylin and eosin staining was performed according to standard procedures. Briefly, sections were dewaxed with xylenes and serial ethanol dilutions then stained with Eosin using the Richard-Allan Scientific Signature Series Eosin-Y package (ThermoScientific).

### Primary motor neuron culture and transfection

Primary motor neurons from e13.5 mouse embryos were isolated, cultured, and transfected by magnetofection as previously described (Fallini et al., 2010; Fallini et al., 2011). For visualization, motor neurons were transfected with GFP to highlight the entire cell. For high resolution imaging, a 60x objective (1.4 NA) was used. Z-series (5 to 10 sections, 0.2 μm thickness) were acquired with an epifluorescence microscope (Ti, Nikon) equipped with a cooled CCD camera (HQ2, Photometrics). Low magnification images were acquired with a 10x or 20x phase objective. Overlapping images of a single cell were reassembled if necessary using Photoshop (Adobe).

## Results

### LSD1 is expressed in neural stem cells *in vivo*

To determine whether LSD1 functions in neural stem cells (NSCs) in mice, we first looked at whether LSD1 is expressed in *Nestin-Cre* positive NSCs *in vivo* in mice. *Nestin-Cre* is strongly expressed in neural tissues beginning at e10.5, during which time NSCs are beginning to proliferate into various progenitor cell populations (Tronche et al., 1999). We performed immunofluorescence for LSD1 and CRE to detect protein levels in the neural tube at embryonic day 12.5 (e12.5). These data show that LSD1 is robustly expressed throughout the neural tube at this time point, including in the *Nestin-Cre* positive NSCs (Fig. 1A-H).

**Figure 1.**
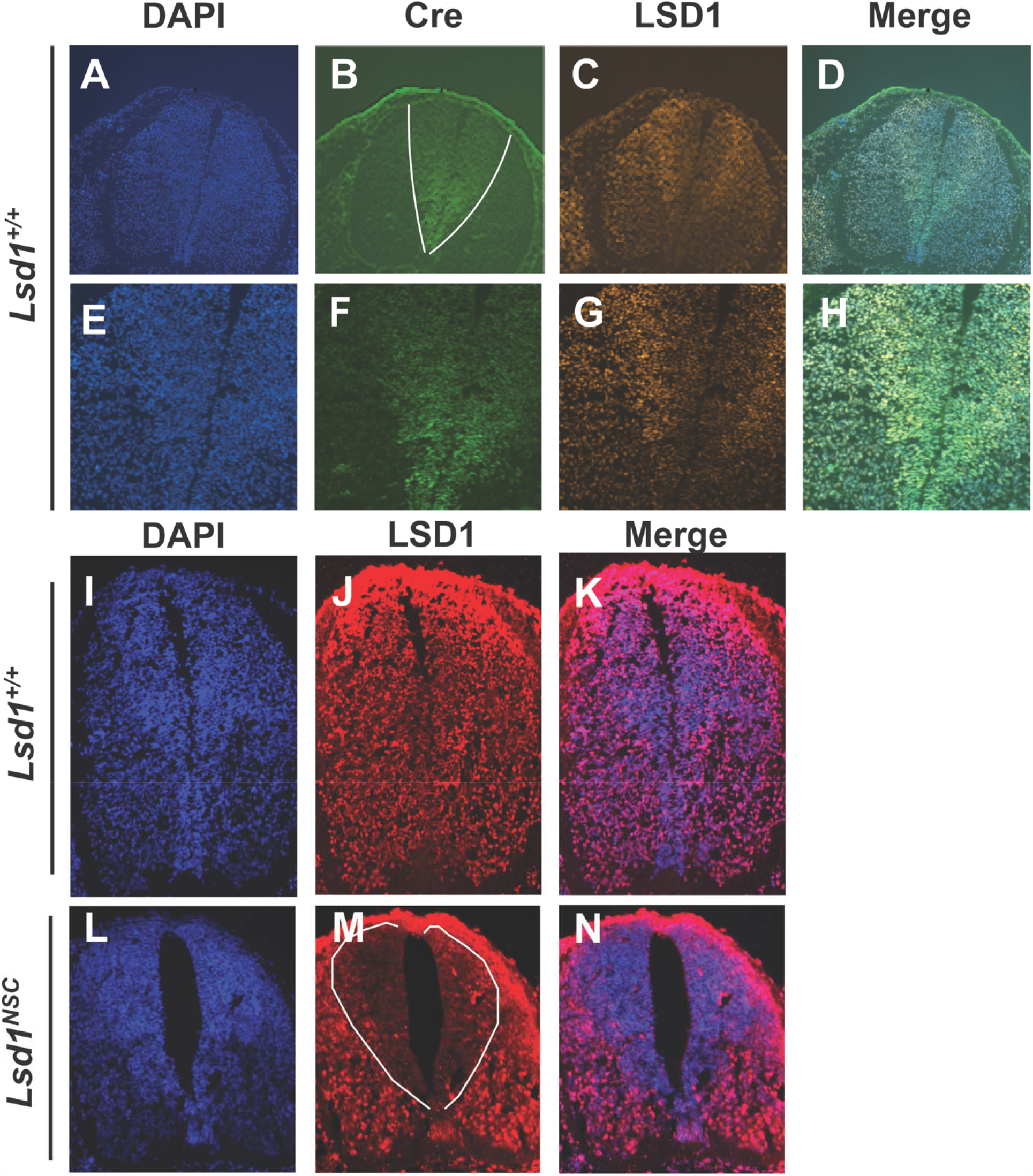
LSD1 is expressed in neural stem cells. Immunofluorescence at e12.5 of *Lsd1*^*+/+*^ control (A-K) and *Lsd1*^*NSC*^ (L-N) embryos, showing DAPI (A,E,I,L), LSD1 (C,G,J,M), Cre (B,F) and Merge (D,H,K,N). CRE+ NSCs outlined in white (B) are magnified in E-H. Expression of LSD1 in these cells is lost in *Lsd1*^*NSC*^ embryos (M).

### *Lsd1*^*NSC*^ animals die postnatally with motor defects

To determine whether there is a phenotype when *Lsd1* is deleted in NSCs, we conditionally deleted *Lsd1* with *Nestin-Cre* (Tronche et al., 1999). We first performed immunofluorescence at e12.5 to confirm that deletion of LSD1 with *Nestin-Cre* results in the loss of LSD1 in NSCs. We found that there was a loss in LSD1 signal in *Lsd1*^*NSC*^ mice compared to controls, suggesting that deletion of LSD1 with *Nestin-Cre* results in the loss of LSD1 in NSCs (Fig. 1I-N). Controls were littermates of *Lsd1*^*NSC*^ animals that were *Cre-*, hereafter referred to as *Lsd1*^*+/+*^ mice.

Next, we examined the survival of *Lsd1*^*NSC*^ mutants compared to controls. From multiple different crosses containing both the *Lsd1* floxed allele and *Nestin-Cre, Lsd1*^*NSC*^ mice were born at expected Mendelian ratios, suggesting there is no embryonic lethality associated with the loss of LSD1 in NSCs (Fig. 2A). In addition, *Lsd1* heterozygotes were observed at the expected frequency at weaning (Fig. 2B). However, by postnatal day 21 (P21), there were zero viable *Lsd1*^*NSC*^ mice observed, compared to the expected frequency of 15.8% which should have yielded 14.6 animals (Fig. 2B). This indicates that 100% of *Lsd1*^*NSC*^ mutants died prior to weaning. Most of *Lsd1*^*NSC*^ animals died between 2-5 days postnatally, with a few animals surviving until 8-11 days.

**Figure 2.**
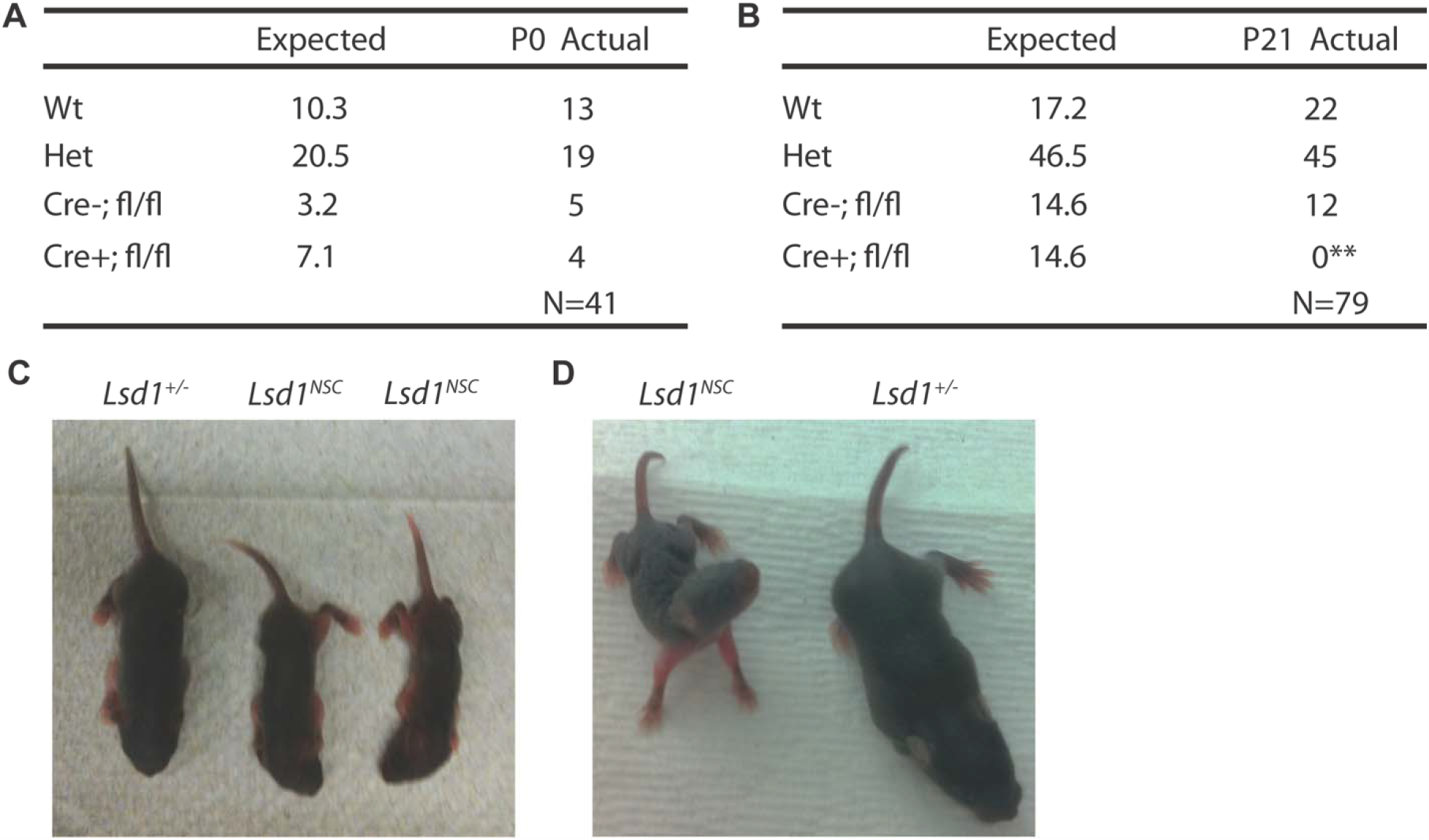
*Lsd1*^*NSC*^ animals do not survive past weaning and show stunted growth. Expected number of mice of each possible genotype (Wt: Wild Type, Het: heterozygous for the *Lsd1* floxed allele with and without Cre, Cre-; fl/fl: homozygous for the *Lsd1* floxed allele without Cre, Cre+; fl/fl: homozygous for the *Lsd1* floxed allele with Cre) versus observed at P0, N=41 from 7 litters (A) and P21, N=79 from 14 litters (B). **Indicates a P<0.01 from chi square analysis. (C,D) Representative images of heterozygous *Lsd1*^*+/-*^ control versus *Lsd1*^*NSC*^ littermates taken at postnatal day 5 and postnatal day 8, with *Lsd1*^*NSC*^ animal engaged in stargazing pose.

To examine the cause of death, we observed *Lsd1*^*NSC*^ mice after birth. *Lsd1*^*NSC*^ animals had severely stunted growth compared to controls (Fig. 2C,D). Additionally, mutants appeared to show difficulty walking steadily, and engaged in stargazing, in which the pup’s head points upward (Fig. 2D, Supplemental Video 1). Taken together, these data show that loss of LSD1 in NSCs results in stunted growth and gross motor defects prior to perinatal lethality.

Due to the gross motor phenotypes observed in *Lsd1*^*NSC*^ mice, we wanted to test whether the initial specification of motor neuron populations embryonically was affected in *Lsd1*^*NSC*^ embryos. To examine this possibility, we performed immunofluorescence at e12.5 using two markers of motor neuron differentiation: Olig2 and HB9. Olig2 marks motor neuron progenitors and HB9 marks motor neurons (Arber et al., 1999; Novitch et al., 2001). At e12.5, there was no difference in the location or quantity of either the Olig-2- or HB9-positive cell populations between *Lsd1*^*NSC*^ mutants and controls, indicating that the initial motor neuron specification was normal (Fig. 3).

**Figure 3.**
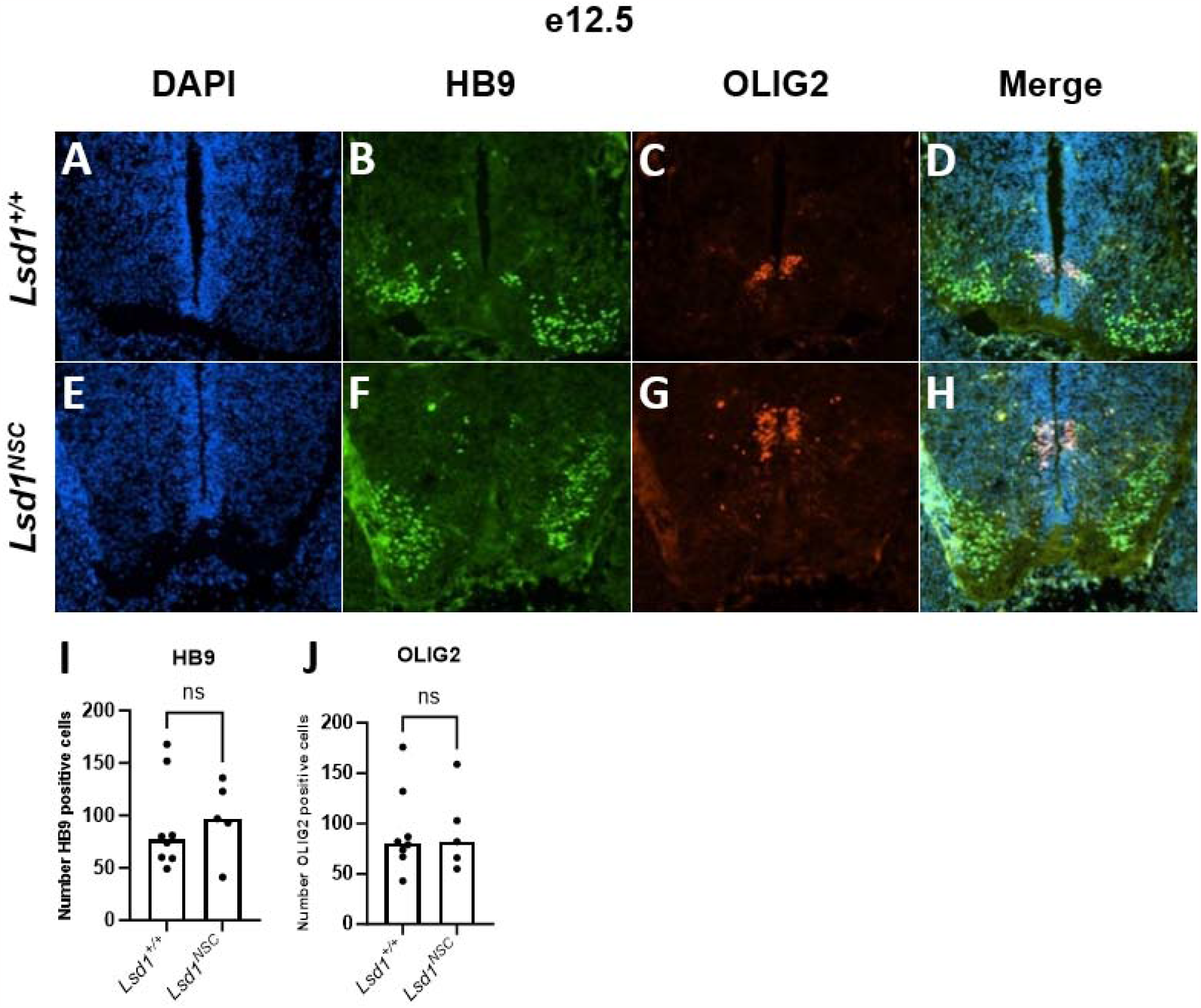
Specification of motor neurons in *Lsd1*^*NSC*^ animals is not affected. Immunofluorescence at e12.5 showing DAPI (A,E), HB9 (B,F) and OLIG2 (C,G) in *Lsd1*^*+/+*^ control (A-D) versus *Lsd1*^*NSC*^ (E-H) embryos, quantified in I and J (*Lsd1*^*+/+*^: N=8, *Lsd1*^*NSC*^: N=5). ns indicates not significant.

### Motor neurons differentiated from LSD1-deficient NSCs inappropriately maintain stem cell proteins

To further investigate whether LSD1-deficient NSCs had defects when differentiating into motor neurons, we cultured motor neurons *in vitro* from e13.5 *Lsd1*^*NSC*^ embryos. Motor neurons derived from *Lsd1*^*NSC*^ appeared morphologically normal (Fig 4. A,B). However, motor neurons derived from *Lsd1*^*NSC*^ embryos continue to inappropriately maintain NSC proteins (Fig. 4C-T).

**Figure 4.**
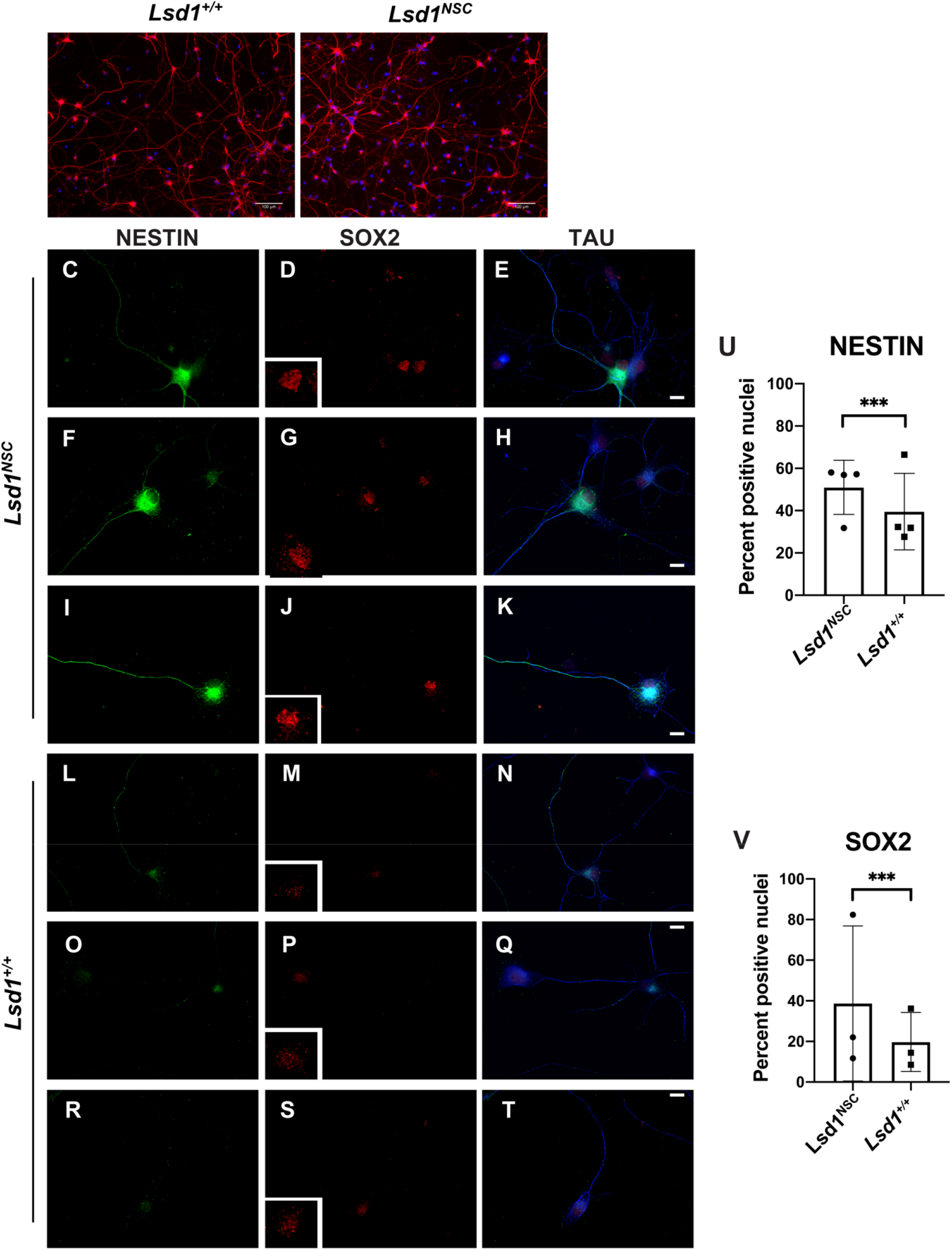
*Lsd1*^*NSC*^ mutant-derived motor neurons inappropriately express stem cell genes. Motor neurons cultured from e13.5 *Lsd1*^*+/+*^ controls (A) and *Lsd1*^*NSC*^ mutants (B) appear morphologically normal. Three examples each of motor neurons cultured from *Lsd1*^*NSC*^ mutants (C-K) and *Lsd1*^*+/+*^ controls (L-T), showing the NSC markers NESTIN (C,F,I,L,O,R) and SOX2 (D,G,J,M,P,S) and the neuronal marker TAU (E,H,K,N,Q,T), quantified in U and V. *** = p<.001, calculated using an unpaired t test.

Normally cultured motor neurons either do not maintain or very weakly maintain the critical NSC proteins NESTIN and SOX2 (Fig. 4L-T). However, upon loss of LSD1 in NSCs, NESTIN and SOX2 continue to persist in differentiated motor neurons (Fig. 4C-K). This resulted in a significant increase in the percentage of nuclei scored as positive for these two NSC proteins compared to controls (Fig. 4U,V). These results suggest that LSD1 is required for the downregulation of NESTIN and SOX2 during motor neuron differentiation.

### Postnatal *Lsd1*^*NSC*^ mutants show abnormal brain morphology *in vivo*

Postnatal *Lsd1*^*NSC*^ animals have gross motor defects. To determine if defects in NSC differentiation give rise to overall defects in brain morphology, we performed hemotoxylin & eosin (H&E) staining on the brains of *Lsd1*^*NSC*^ animals that died perinatally. Compared to controls the hippocampus and cerebellum of *Lsd1*^*NSC*^ mutants looked disorganized (Fig. 5). Also, the overall size of the cerebellum in *Lsd1*^*NSC*^ mice was much smaller than controls (Fig. 5C,D). In addition to the overall changes in morphology, we observe a large number of nuclei in the *Lsd1*^*NSC*^ hippocampus that appear to be smaller and more pyknotic than in controls (Fig. 5E,F). Overall, these data suggest that loss of LSD1 in NSC leads large changes in brain morphology and structure.

**Figure 5.**
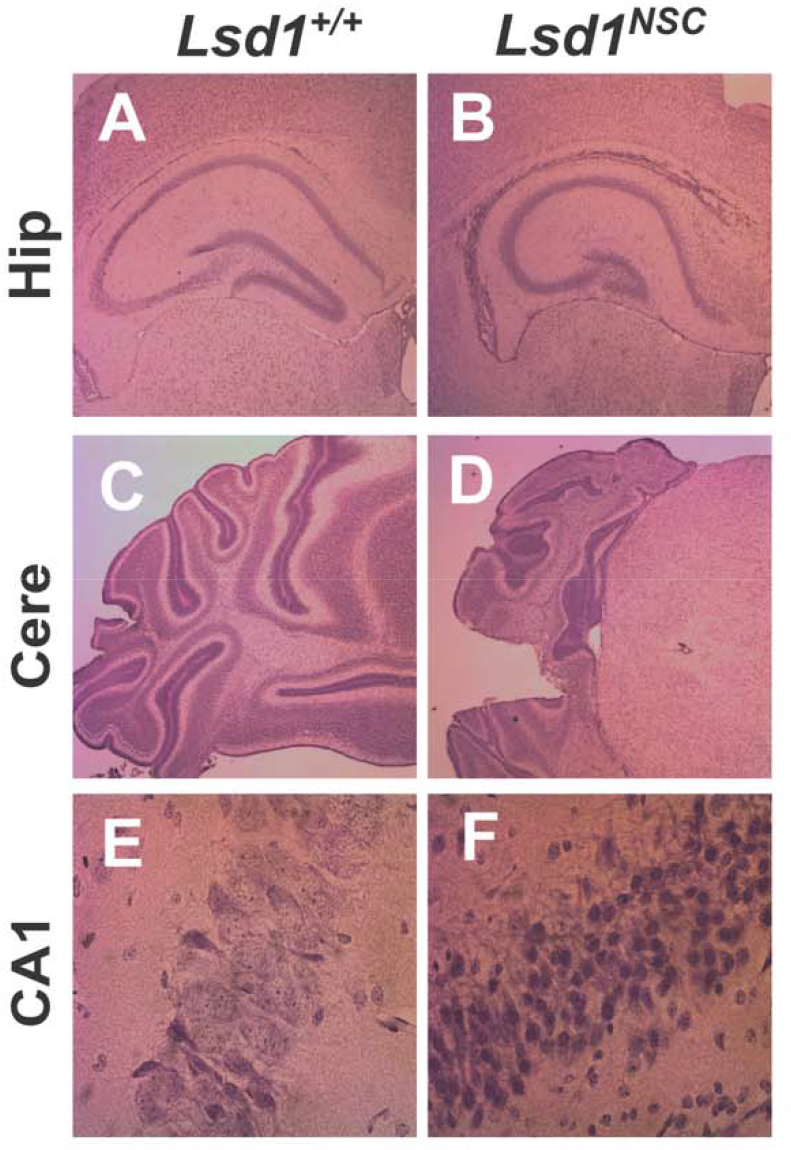
*Lsd1*^*NSC*^ mutant animals show brain morphology defects *in vivo*. H&E histology of (A,B) hippocampus (Hip), (C,D) cerebellum (Cere) and (E,F) CA1 region of the hippocampus (CA1) from *Lsd1*^*+/+*^ control (A,C,E) versus *Lsd1*^*NSC*^ (B,D,F) postnatal animals.

## Discussion

Consistent with previous data showing that LSD1 is expressed in adult NSC in the mouse brain (Sun et al., 2010), we found that LSD1 is expressed in mouse embryonic NSCs. Therefore, to determine whether LSD1 functions in NSC differentiation *in vivo* in mice, we conditionally deleted *Lsd1* in NSCs using *Nestin-Cre*. We find that loss of LSD1 in *Nestin* positive NSCs results in 100% perinatal lethality. This suggests that LSD1 is essential for the proper establishment of the nervous system. Most of the *Lsd1*^*NSC*^ animals survived beyond 24 hours and grew. This indicates that *Lsd1*^*NSC*^ mutants can breathe and feed. However, *Lsd1*^*NSC*^ animals have severely stunted growth, indicating a defect in obtaining proper nutrition. In addition, *Lsd1*^*NSC*^ animals have an obvious motor coordination defect, suggesting that LSD1 is required in NSCs for the proper functioning of the neuromuscular system.

To investigate the underlying causes of the motor coordination defect, we determined whether motor neurons are being properly specified from the NSC population when LSD1 is eliminated in NSCs. Surprisingly, we found that LSD1 is dispensable for the proper specification of the Olig2-positive motor neuron progenitors and the initial HB9-positive motor neuron population *in vivo*. This result differs from other stem cell populations in the mouse, such as hematopoietic stem cells, testis stem cells, naïve B cells, trophoblast stem cells and satellite SCs, where LSD1 is essential for the proper initial differentiation (Haines et al., 2018; Kerenyi et al., 2013; Lambrot et al., 2015; Myrick et al., 2017; Saleque et al., 2007; Tosic et al., 2018). This hints that the function of LSD1 in NSCs may subtly differ from other stem cell populations.

Nevertheless, we find that when LSD1 is not present in NSCs, critical NSC proteins (NESTIN and SOX2), continue to persist abnormally in differentiated neurons. This suggests that LSD1 is required to downregulate these critical NSC proteins for proper NSC differentiation. Previously, LSD1 has been shown to decommission enhancers at critical stem cell genes during stem cell differentiation in the mouse by removing H3K4me1/2 (Kerenyi et al., 2013; Lambrot et al., 2015; Myrick et al., 2017; Whyte et al., 2012). Our data are consistent with the possibility that LSD1 functions similarly in NSC differentiation.

It was surprising that LSD1-deficient NSCs can differentiation into morphologically normal motor neurons from NSCs despite the persistence of critical NSC proteins. This indicates one potential way in which the differentiation of NSCs may differ from the differentiation of other mouse stem cell populations. It is possible that in NSCs the down regulation of the SC program may not be coupled to the activation of the differentiation program. Whereas in other stem cell populations, the down regulation of the SC program may be essential for the proper activation of the differentiation program.

Despite our finding that motor neurons can be made from NSCs without LSD1, we observe perinatally lethality and severe motor coordination defects in *Lsd1*^*NSC*^ pups. Thus, it is possible that motor neurons may be specified but do not function properly in these mice. In this case, the continued inappropriate maintenance of the SC program may interfere with the function of motor neurons. Importantly, if the inappropriate maintenance of the SC program can interfere with the ongoing function of motor neurons, it might be possible to reverse these defects by shutting off the SC program.

*Lsd1*^*NSC*^ pups also have severe defects in brain morphology. Thus, it is possible that neurons are initially made but are then dying due to loss of LSD1. LSD1 has previously been shown to be continually required for the survival of hippocampus and cortex neurons. When LSD1 is lost, these neurons become pyknotic and die (Christopher et al., 2017). Consistent with this possibility, we observe pyknotic nuclei in the hippocampus of the *Lsd1*^*NSC*^ pups. This raises the possibility that these neurons may be degenerating perinatally.

The data presented here provide the first evidence *in vivo* that LSD1 is a critical regulator of NSC differentiation in mice. As a result, the *Lsd1*^*NSC*^ mice will provide an excellent *in vivo* model for understanding how the inappropriate maintenance of SC program interferes with neuronal function. In addition, these findings add to the growing body of literature showing that LSD1 is essential for stem cell differentiation.

## Acknowledgements

We are grateful to members of the Katz lab, as well as T. Caspary, R. Bear, and C. Bean for their helpful discussion, critical reading of the manuscript and for strains.

## Funding

This work was funded by grants to D.J.K. (NSF IOS1931697, Emory University Research Council, pilot funding from P01GM085354 (Dalton, PI) and NINDS 1R56NS122964-01A1). A.S. was supported by NIH F31 5F31HD098816. M.R. and A.S. were supported by the Emory GMB training grants (T32GM008490-21 and T32GM149422-01). This study was supported in part by the Mouse Transgenic and Gene Targeting Core (TMF), which is subsidized by the Emory University School of Medicine and is one of the Emory Integrated Core Facilities. Additional support was provided by the National Center for Advancing Translational Science of the National Institutes of Health (UL1TR000454).

## Author Contributions

E.F., M.R., A.S., D.M and D.J.K. conceived and designed the study. A.S., E.F., M.R. and D.J.K wrote the manuscript. E.F., M.R., A.S., and D.M. performed experiments, analyzed data and interpreted results under the direction of D.J.K.. C.F. cultured and performed immunofluorescence on the motor neurons under the direction of G.B.. All authors discussed the results.

## Notes

### Competing Interest Statement

The authors have declared no competing interest.

